# SPAK as a candidate mediator of TRPV4-induced NKCC1 activation and a possible pharmacological entry point in posthemorrhagic hydrocephalus

**DOI:** 10.64898/2026.07.26.740775

**Authors:** Trine L. Toft-Bertelsen, Nanna MacAulay

## Abstract

Hydrocephalus arises from pathological disturbances in cerebrospinal fluid (CSF) homeostasis, yet current treatment relies almost exclusively on neurosurgical diversion procedures that frequently require surgical revision. No specific and efficient pharmacological alternative is available as a complement to the invasive neurosurgery due to our lack of understanding of the molecular regulators of CSF secretion. By *in vivo* determination of CSF dynamics in rats and *in vitro* quantification of choroid plexus transporter activity, we demonstrate that the STE20-proline-alanine-rich kinase (SPAK) is a critical modulator of CSF secretion via its regulation of the Na^+^/K^+^/2Cl^-^ cotransporter 1 (NKCC1) and the Na⁺/K⁺-ATPase. Systemic administration of a SPAK inhibitor after a mimicked hemorrhagic event attenuated posthemorrhagic hydrocephalus formation 24h post-hemorrhage. Activation of the choroid plexus transient receptor potential vanilloid 4 (TRPV4) ion channel induced hydrocephalus through CSF hypersecretion. This TRPV4-mediated hypersecretion occurred via activation of NKCC1, not the Na^+^/K^+^-ATPase, and required SPAK activity as a molecular link. Together, these findings identify SPAK as a central integrator of TRPV4-dependent signaling and choroid plexus transporter activity, positioning the TRPV4-SPAK axis as a potential pharmacological target for modulating CSF dynamics in hydrocephalus and other pressure-related pathologies.

## Introduction

Hydrocephalus is a neurological disorder characterized by the abnormal accumulation of cerebrospinal fluid (CSF) in the ventricular system, resulting in increased intracranial pressure (ICP) and potential cerebral damage [1, 2]. Traditionally, hydrocephalus is treated with surgical intervention to divert excessive CSF from the ventricles and into the abdominal cavity through a ventricular shunt implantation or through external ventricular drainage [3, 4]. Such surgery, while effective, is frequently associated with complications such as infection and/or shunt malfunctions that require semi-acute shunt revision [5, 6]. Alternative pharmacological approaches have so far proven suboptimal [7–9]. An understanding of the molecular mechanisms underlying CSF production and their regulation in physiology and pathophysiology could therefore pave the way for pharmacological therapies as a supplement or alternative to invasive neurosurgery.

The choroid plexus is a specialized tissue residing in the brain ventricles and responsible for the majority of CSF production [10, 11]. Recent research has highlighted the role of the STE20-proline-alanine-rich kinase (SPAK), which is highly expressed in choroid plexus [12] and an established cell volume sensor [13], as a key instigator of the CSF hypersecretion that appears to be a component of acquired hydrocephalus caused by brain infection or following a hemorrhagic event [14–16]. Anti-inflammatory strategies, and pharmacological or genetic inhibition of SPAK, attenuated the CSF hypersecretion in animal models of these pathologies and thus reduced the hydrocephalus formation [14, 15].

The transient receptor potential vanilloid 4 (TRPV4) ion channel is a mechanosensitive ion channel with a well-established cell volume-sensing modality [17–19] and highly expressed on the luminal membrane of the choroid plexus [9, 12, 16]. Intraventricular activation of TRPV4 increases the rate of CSF secretion in healthy rats [16] and in a human choroid plexus papilloma cell line [20], whereas i.p. delivery of the TRPV4 inhibitor RN1734 fails to reach its luminal target and affect the CSF secretion rate [21]. Pharmacological inhibition of TRPV4, accordingly, reduces hydrocephalus formation in both a rodent model of congenital hydrocephalus [22] and in one of post-hemorrhagic hydrocephalus [16]. TRPV4 is directly and acutely activated by the plasma lipid lysophospahtic acid (LPA) that binds to the ion channel [23] upon entry into the ventricles with the hemorrhagic event in both humans and rats, and which can induce hydrocephalus following a stand-alone intraventricular bolus delivery [16]. The LPA-induced CSF hypersecretion appear to occur, at least in part, via activation of the Na^+^,K^+^,2Cl^-^ cotransporter 1 (NKCC1), a key contributor to CSF secretion in rats, mice, and dogs [15, 24–26], a pattern similar to that obtained with a pharmacological activator of TRPV4 [16]. The TRPV4-mediated activation of NKCC1 is blunted upon SPAK inhibition [16], suggesting that TRPV4-mediated control of CSF secretion could act via SPAK. SPAK may, as such, act as a convergence hub integrating inflammatory, osmotic, mechanosensitive, and hemorrhage-driven signals into a common CSF secretory response in health and disease.

Here, we reveal the SPAK-dependent regulation of key components in the CSF-secreting molecular machinery of healthy rats and reveal the crosstalk between TRPV4 signaling and the SPAK-NKCC1 axis as a critical regulatory mechanism underlying a component of post-hemorrhagic hydrocephalus formation in rats.

## Results

### SPAK regulates CSF production by modulating choroid plexus transporter activity

To reveal SPAK-mediated regulation of the CSF production, we used an *in vivo* imaging strategy that allows imaging of the ventricular system in anesthetized rats immediately after intracerebroventricular delivery of a fluorescent dye as a proxy of CSF production. This proxy measurement has been demonstrated to match that obtained with both the ventricular cisternal perfusion assay and that of the ‘direct method’ [16, 24, 25, 27]. This experimental strategy is thus considered a validated quantification of *relative* changes in CSF secretion rates upon inhibitor administration, and will be referred to henceforth as CSF secretion. The dye flow is visualised as superimposed pseudo-colour fluorescence, with the dye intensity quantified in the region of interest, placed in line with the skull landmark lambda on a white light image (**Fig. 1A**), in which the fluorescence intensity was recorded every 30 sec over a period of 3 min (**Fig. 1Ai**). Post-mortem verification of correct targeted dye delivery was demonstrated post-recording (**Fig. 1Aii)**. The fluorescence intensity recorded over time was normalized to the intensity of the first image (**Fig. 1B**). Quantification of the dye flow rate demonstrated a reduced CSF secretion rate upon SPAK inhibition with ZT-1a (intraperitoneal delivery 30 minutes before recording; 0.13 ± 0.01 min^-1^ in control rats and 0.05 ± 0.02 min^-1^ in ZT-1a - treated rats, n = 6, **Fig. 1C**, P < 0.001, See **Suppl. Fig. S1** for the collected data set). To identify the molecular coupling between SPAK activity and the CSF secretion machinery, we performed isotope flux assays on acutely isolated rat choroid plexus to determine the SPAK-dependent activity of choroid plexus transporters involved in CSF secretion, the Na^+^/K^+^-ATPase and the NKCC1 (illustrated as an inset in **Fig. 1D**). ^86^Rb^+^ acts as a congener for K^+^ and can, as such, replace K^+^ in its binding sites in both the Na^+^/K^+^-ATPase and the NKCC1, where the ^86^Rb^+^ transport rate represents a read-out of transporter activity. The Na^+^/K^+^-ATPase activity was determined with an ^86^Rb^+^ uptake assay in the absence and presence of the Na^+^/K^+^-ATPase inhibitor ouabain and/or ZT-1a, both of which reduced the ^86^Rb^+^ uptake (14.1 ± 0.5 cpm in control, 2.2 ± 0.2 cpm with ouabain exposure, 3.8 ± 0.3 cpm with ZT-1a exposure, and 1.2 ± 0.3 cpm with ouabain and ZT-1a exposure, n = 5 of each, **Fig. 1D**). Although there is residual background uptake presumably from other transporters and channels, the ouabain-sensitive fraction represents the Na^+^/K^+^-ATPase-mediated activity, which was significantly reduced upon SPAK inhibition (11.9 ± 0.6 cpm in control vs 2.5 ± 0.7 cpm with ZT-1a, n = 5 of each, **Fig. 1E**, P < 0.001). The outwardly directed NKCC1-activity was determined with an ^86^Rb^+^ efflux assay as a function of time in the absence and presence of the NKCC1 inhibitor bumetanide and/or the SPAK inhibitor ZT-1a (**Fig. 1F**). The ^86^Rb^+^ efflux rate (0.39 ± 0.03 min^−1^in control, n = 6) was robustly inhibited by bumetanide (0.11 ± 0.03 min^−1^, n = 6, P < 0.001) and ZT-1a (0.12 ± 0.04 min^−1^, n = 6, P < 0.001, **Fig. 1G**). The ^86^Rb^+^ efflux rate was similar (P = 0.99) in the presence of both inhibitors, suggesting that NKCC1 is inactive when SPAK activity is abolished. Taken together, these data indicate that SPAK activity is required for basal choroid plexus transporter activity and, consequently, the transporter-mediated contribution to the CSF production.

**Fig. 1.**
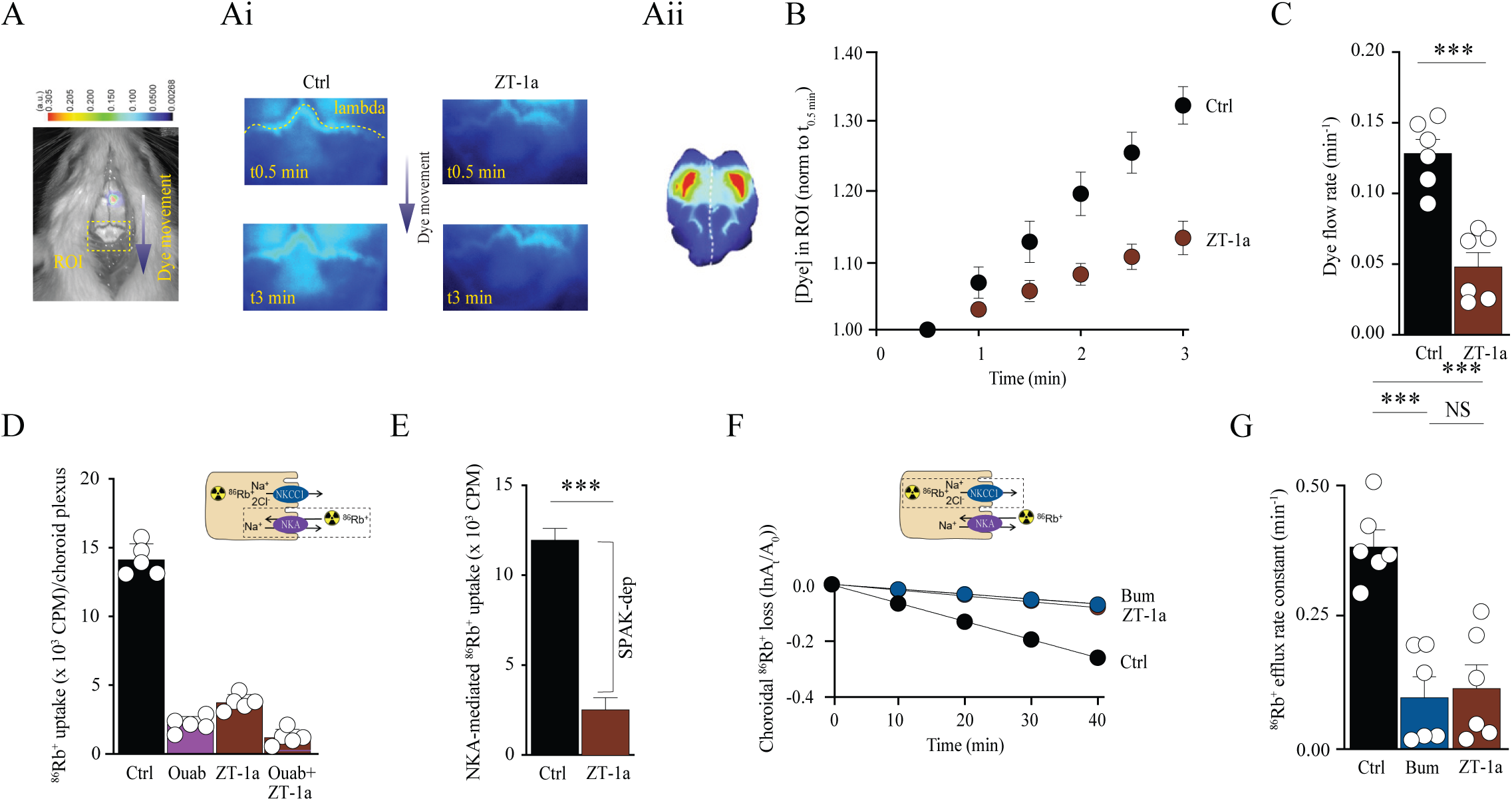
SPAK regulates CSF production by modulating choroid plexus transporter activity. **A** Representative image of a rat after injection of IRDye 800CW carboxylate dye (superimposed pseudo-color). The square placed in line with lambda indicates the area of dye content quantification as it moves rostrally (**Ai**), and a rat brain showing correctly targeted dye delivery in mid-sagittal sections (**Aii**). **C** Quantification of the dye intensity (as a proxy for CSF production) determined from linear regression in **B** over a 3-minute time window with (n = 6) or without (n = 6) SPAK inhibition by ZT-1a delivered intraperitoneally. **D** ^86^Rb^+^ influx (measured in counts per minute; cpm) in the presence or absence of ouabain and/or ZT-1a (n = 5 in each group). Inset: Schematic illustration of the principle behind the ^86^Rb^+^ isotope flux assays. **E** Na^+^/K^+^-ATPase (NKA)-mediated ^86^Rb^+^influx per choroid plexus (ouabain-sensitive fractions; n = 5 in both groups). **F** Loss of ^86^Rb^+^ from the choroid plexus as a function of time in the presence or absence of bumetanide (Bum) or ZT-1a. The y-axis is the natural logarithm of the choroid plexus ^86^Rb^+^ amount left at time T divided by the initial amount at time 0 (A0) [16]. **G** Efflux rate constant for ^86^Rb^+^ in the presence (n = 6) or absence (n = 6) of bumetanide or ZT-1a (n = 6). See Suppl. Fig. S1 for the collected data set. Statistical evaluation with *Student’s* t-test (panel C and E) or one-way ANOVA with Tukey’s post-hoc test (panel G). ***P < 0.001, NS: not significant.

### SPAK-inhibition alters CSF dynamics following brain hemorrhage

To determine the SPAK-dependent modulation of the brain CSF dynamics in a pathological setting, we introduced ZT-1a in a rat model of intraventricular hemorrhage (IVH) that develops posthemorrhagic hydrocephalus within 24 h after delivery of autologous blood into the right lateral ventricle of anesthetized rats. The intraventricular hemorrhage-induced ventricular enlargement was evident with MRI (see representative 3D-volumetry of the ventricular system in **Fig. 2A**), where the lateral ventricular volume was significantly elevated in the IVH rats with the inflicted hemorrhagic insult (57.7 ± 6.0 mm^3^, n = 8) compared to that of control rats undergoing sham surgery with intraventricular delivery of saline (7.07 ± 1.8 mm^3^, n = 6, P < 0.001, **Fig. 2B**). IVH rats treated with ZT-1a 2 h post-IVH induction (re-dosed at 12 h) presented with significantly smaller ventricular volumes (37.4 mm^3^ ± 5.7, n = 8) than those of the non-treated IVH rats (P < 0.05), although still significantly larger than those of the control rats (P < 0.01, **Fig. 2B**, see representative MRI scans in **Suppl. Fig. S5A**). This pattern was also observed for the total CSF pool, with no difference in body weight as a function of treatment (**Suppl. Fig. S2A-B**). These findings were mimicked on the total brain water content, which increased in rats with IVH compared to that observed in sham-operated control rats (from 3.64 ± 0.04 mL brain H2O/g dry brain, n = 7 to 4.58 ± 0.18 mL brain H2O/g dry brain n = 8 in IVH rats, P < 0.001, **Fig. 2C**). The IVH-induced brain water elevation was significantly reduced by 2 h post-IVH induction administration (re-dosed at 12 h) of the SPAK inhibitor ZT-1a (3.98 ± 0.11 mL brain H2O/g dry brain, n = 8, P < 0.05) to a level not significantly different from control, P = 0.17, **Fig. 2C**. These results indicate that inhibition of SPAK in a setting of brain hemorrhage reduces the ensuing posthemorrhagic hydrocephalus development in the acute phase.

**Fig. 2.**
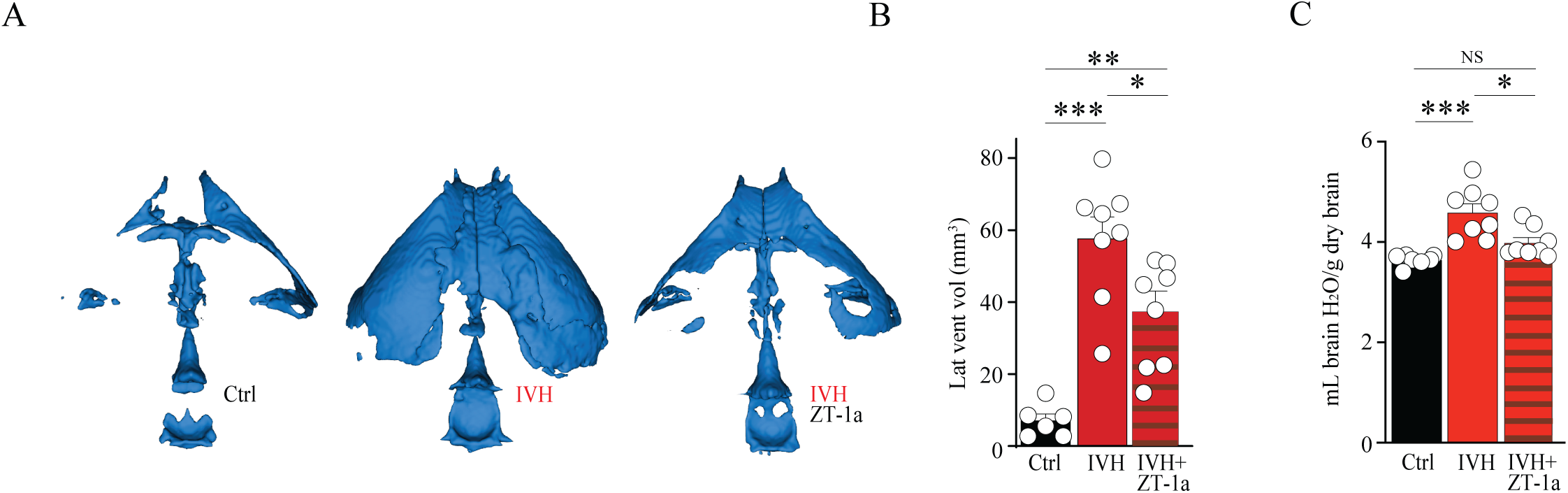
SPAK-inhibition alters CSF dynamics following brain hemorrhage. **A** Representative 3D projections of a control rat, an intraventricular hemorrhage (IVH) rat and an IVH rat treated with ZT-1a, demonstrating the ventricular volume 24 h post-insult. **B** Lateral ventricle volumes of control rats (n = 6), IVH rats (n = 8), and IVH rats treated with ZT-1a (n = 8) quantified from MRI, see **Suppl. Fig. S5A** for representative MRI scans. **C** Brain water content quantified from control rats, IVH rats and IVH rats treated with ZT-1a. Statistical evaluation with one-way ANOVA with Tukey’s post-hoc test. *P < 0.05, **P < 0.01, ***P < 0.001, NS: not significant.

### TRPV4-mediated modulation of CSF dynamics

Previous observations suggest that a SPAK-mediated modulation of the CSF dynamics could, at least in part, originate from acute upstream TRPV4 activity [16]. To resolve sustained TRPV4-mediated effects on CSF homeostasis, we administered the TRPV4 activator GSK1016790A (henceforth GSK101) intracerebroventricularly in anaesthetised rats with subsequent quantification of the brain water content 24 h after administration. The total brain water content was increased by GSK101 administration (4.27 ± 0.27 mL brain H2O/g dry brain, n = 5) compared to that obtained in control (vehicle-treated) rats (3.68 ± 0.06 mL brain H2O/g dry brain, n = 5, P < 0.05, **Fig. 3A**; note that per oral and intraperitoneal administration did not induce increased brain water, possibly due GSK101 not crossing the blood-CSF barrier in sufficient degree to target the luminally expressed TRPV4, see **Suppl. Fig. S3A-B**). The elevated brain water content may arise, at least in part, by increased CSF secretion, as intracerebroventricular delivery of the TRPV4 activator GSK101 acutely increased the CSF secretion rate by 40% (0.13 ± 0.01 min^-1^, n = 6 in control rats vs. 0.18 ± 0.01 min^-1^, n = 6 in TRPV4-activated animals, P < 0.01, **Fig. 3B-C**), confirming that TRPV4 activation by GSK101 regulates CSF secretion in healthy rats.

**Fig. 3.**
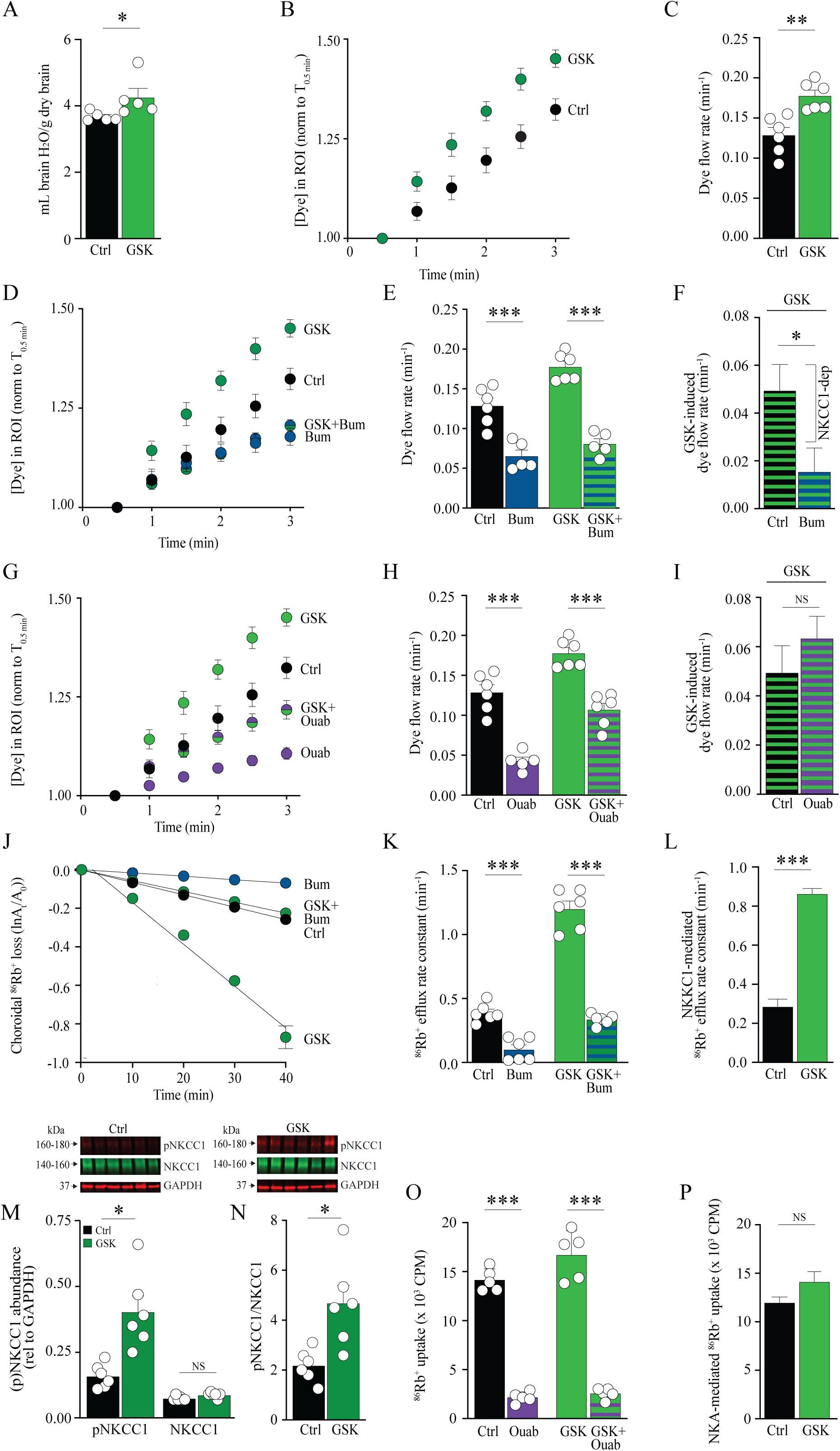
TRPV4-mediated modulation of CSF dynamics associated with NKCC1, not Na^+^/K^+^-ATPase, hyperactivity. **A** Brain water content quantified from control rats (n = 5) and rats treated with GSK101 intracerebroventricularly (n = 5). **C** Quantification of the dye intensity in the region of interest (as a proxy for CSF production) determined from linear regression in **B** over the 3-minute time window from control (n = 6) and GSK101 delivered intracerebroventricularly (n = 6). **E+H** Quantification of the dye intensity determined from linear regression in **D+G** over the 3-minute time window from control (n = 6), GSK101 delivered intracerebroventricularly (n = 6), bumetanide (Bum) (n = 5) or GSK101+Bum (n = 5), or ouabain (Ouab) (n = 5) or GSK101+Ouab (n = 6), respectively. **F+I** GSK101-induced dye flow rates upon NKCC1 or Na^+^/K^+^-ATPase inhibition, respectively. **J** Loss of ^86^Rb^+^ from the choroid plexus as a function of time in the presence or absence of bumetanide (Bum) or GSK101. The y-axis is the natural logarithm of the choroid plexus ^86^Rb^+^ amount left at time T divided by the initial amount at time 0 (A0) [16]. **K** Efflux rate constant for ^86^Rb^+^ in the presence (n = 6) or absence (n = 6) of bumetanide, GSK101 (n = 6) or GSK101+Bum (n = 6). **L** NKCC1-mediated ^86^Rb^+^ efflux rate constant (bumetanide-sensitive fractions). **M** NKCC1 and pNKCC1 abundance (relative to GAPDH loading control) in excised choroid plexus lysates after GSK101-treatment and western blotting (n = 6 biological replicates). **N** Ratio of pNKCC1/NKCC1) from paired samples in panel M (n = 6). **Insets** Representative fractions of the western blot. Insets illustrate representative fractios of the western blot, see Suppl. Fig. S4A for the complete uncropped membrane. **O** ^86^Rb^+^ influx (measured in counts per minute; cpm) in the presence (n = 5) or absence (n = 5) of ouabain and/or GSK101 (n = 5). **P** Na^+^/K^+^-ATPase (NKA)-mediated ^86^Rb^+^influx per choroid plexus (ouabain-sensitive fractions). See Suppl. Fig. S1 for the collected data set. Statistical evaluation with *Student’s* t-test (panel C, F, I, L, N, P) or one-way ANOVA with Tukey’s post-hoc test. *P < 0.05, **P < 0.01, ***P < 0.001, NS: not significant.

### GSK101-induced CSF hypersecretion associates with NKCC1, not Na^+^/K^+^-ATPase, hyperactivity

To resolve the molecular origin of the TRPV4-induced CSF hypersecretion of brain water in healthy rats, we determined the CSF secretion rate with inclusion of inhibitors of the key choroid plexus transporters, the NKCC1 and the Na^+^/K^+^-ATPase. Intracerebroventricular delivery of the NKCC1 inhibitor bumetanide reduced the CSF secretion rate approximately 50% (0.07 ± 0.01 min^-1^, n=5) compared to control animals (0.13 ± 0.01 min^-1^, n = 6, P < 0.001, **Fig. 3D-E**), as earlier demonstrated [25]. To determine the GSK101 activation of the NKCC1-mediated contribution to the CSF secretion, we compared the GSK101-mediated increase in CSF secretion in the absence and presence of the NKCC1 inhibitor bumetanide **(Fig. 3D-E)**. With active NKCC1, GSK101 increased the CSF secretion rate by 0.05 ± 0.01 min^-1^ (0.18 ± 0.01 min^-1^, n = 6 in GSK101-treated rats vs 0.13 ± 0.01 min^-1^, n = 6 in control rats), which was significantly reduced to 0.02 ± 0.01 min^-1^, n = 5, P < 0.05 (**Fig. 3F)** upon NKCC1 inhibition during the GSK101 administration (0.08 ± 0.01 min^-1^ in GSK101+bumetanide-treated rats vs 0.07 ± 0.01 min^-1^ in bumetanide-treated rats, n = 5, **Fig. 3E**), suggesting that the TRPV4-mediated CSF hypersecretion requires NKCC1 activity.

The inclusion of ouabain, an inhibitor of the Na^+^/K^+^-ATPase, reduced the CSF secretion (0.04 ± 0.01 min^-1^, n = 6) compared to control rats (0.13 ± 0.01 min^-1^, n = 6, P < 0.001, **Fig. 3G-H**), as previously demonstrated [25]. To determine GSK101 activation of the Na^+^/K^+^-ATPase-mediated contribution to the CSF secretion, we compared the GSK101-mediated increase in CSF secretion in the absence and presence of the Na^+^/K^+^-ATPase inhibitor **(Fig. 3G-H)**. With active Na^+^/K^+^-ATPase, GSK101 increased the CSF secretion rate by 0.05 ± 0.01 min^-1^ (0.18 ± 0.01 min^-1^, n = 6 in GSK101-treated rats vs 0.13 ± 0.01 min^-1^, n = 6 in control rats), which was similar to the GSK101-mediated increase in CSF secretion when the Na^+^/K^+^-ATPase was inhibited during the GSK101 administration (0.06 ± 0.01 min^-1^, n = 6 vs. 0.05 ± 0.01 min^-1^, n = 6, P = 0.35, **Fig. 3I),** suggesting that the TRPV4-mediated CSF hypersecretion is not mediated by TRPV4-induced activation of the Na^+^/K^+^-ATPase.

To determine whether the dependency of NKCC1, but not the Na^+^/K^+^-ATPase, in GSK101-induced hypersecretion was reflected in choroidal transport activity, we quantified their transport activity in acutely isolated choroid plexus with isotope flux assays in the absence and presence of GSK101. The NKCC1-mediated fraction of the ^86^Rb^+^ efflux rate (0.29 ± 0.04 min^−1^) was obtained as the difference between the ^86^Rb^+^ efflux rate in the control setting (0.39 ± 0.03 min^−1^) and that in the presence of bumetanide (0.10 ± 0.03 min^−1^, n = 6 of each, **Fig. 3J-L**). The NKCC1-mediated ^86^Rb^+^ efflux rate (0.29 ± 0.04 min^-1^ black column in Fig. 3L) was tripled in the presence of GSK101 (0.86 ± 0.06 min^−1^, n = 6, P < 0.001, **Fig. 3L**), supporting a direct TRPV4-mediated activation of NKCC1. However, the small GSK-induced, bumetanide-insensitive component of 0.23 ± 0.02 min^−1^, n = 6 observed in **Fig. 3K** (GSK+BUM subtracted BUM), suggests an additional moderate non-NKCC1-mediated K+ efflux route activated by TRPV4. Given the well-established phosphorylation-dependent activation of NKCC1, we confirmed a TRPV4-dependent increase in NKCC1 phosphorylation status by immunoblotting excised choroid plexus tissue following ex vivo GSK101 treatment (pairwise comparison between the two choroid plexuses from the lateral ventricle in a given rat, **Fig. 3M-N + insets)**. The phosphorylated NKCC1 abundance (pNKCC1, ratio to GAPDH loading control) increased from 0.16 ± 0.02 in control to 0.41 ± 0.06 following TRPV4 activation, n = 6, P < 0.05, **Fig. 3M**, with no change in overall NKCC1 abundance (P > 0.99). The ratio of pNKCC1/NKCC1 was thus significantly increased following GSK exposure (2.18 ± 0.26 in control vs 4.67 ± 0.71 following TRPV4 activation, n = 6, P < 0.05, **Fig. 3N**), see **Suppl. Fig. S4A** for the complete, uncropped blot. The Na^+^/K^+^-ATPase-mediated fraction of the ^86^Rb^+^ influx (11.9 ± 0.8 cpm, n = 6, **Fig. 3P**) was obtained as the difference between the ^86^Rb^+^ influx in the control setting (14.1 ± 0.5 cpm, n = 6) and that in the presence of ouabain (2.2 ± 0.2, n = 6, P < 0.001, **Fig. 3O**). This Na^+^/K^+^-ATPase-mediated ^86^Rb^+^ influx was similar in the presence of GSK101 (14.1 ± 1.3 cpm, n = 6, P = 0.35, **Fig. 3P**), suggesting that TRPV4 activation does not influence the Na^+^/K^+^-ATPase transport activity.

### GSK101-induced CSF hyperdynamics depends on SPAK

To reveal the SPAK-dependency of the TRPV4-mediated CSF dynamics, we performed ventricular volumetry on rats following 24 h intracerebroventricular treatment with GSK101 with and without the SPAK inhibitor ZT-1a (see representative MRI scans in **Suppl. Fig. S5B** and 3D-volumetry of the lateral, third and fourth ventricles in **Fig. 4A**). TRPV4 activation increased the lateral ventricle volume 3-fold (from 7.53 ± 0.83 mm^3^ in control rats to 27.3 ± 3.5 mm^3^ in GSK101-treated rats, n = 7, P < 0.001, **Fig. 4B**), demonstrating that TRPV4-activation in itself can induce ventriculomegaly in healthy rats. Inclusion of the SPAK inhibitor ZT-1a (administered 30 min pre-delivery of GSK101) reduced the GSK101-induced ventriculomegaly to a mere 12.9 ± 2.3 mm^3^, n = 7, P < 0.01, which was not significantly different from the control group, P = 0.83, **Fig. 4B**. These data demonstrate a SPAK-dependency of the TRPV4-mediated ventriculomegaly. The findings were mimicked on the total brain water content, which increased in GSK101-treated rats (4.28 ± 0.19 mL brain H2O/g dry brain, n = 7) compared to control rats (3.63 ± 0.03 mL brain H2O/g dry brain, n = 7, P < 0.01, **Fig. 4C**). Inclusion of the SPAK inhibitor ZT-1a reduced the GSK101-induced brain water accumulation (3.84 ± 0.09 mL brain H2O/g dry brain, n = 7, P < 0.05) to the level of that of the control group, P = 0.53, **Fig. 4C**. These data indicate that the TRPV4-mediated CSF accumulation occurs via a SPAK-controlled signalling pathway.

**Fig. 4.**
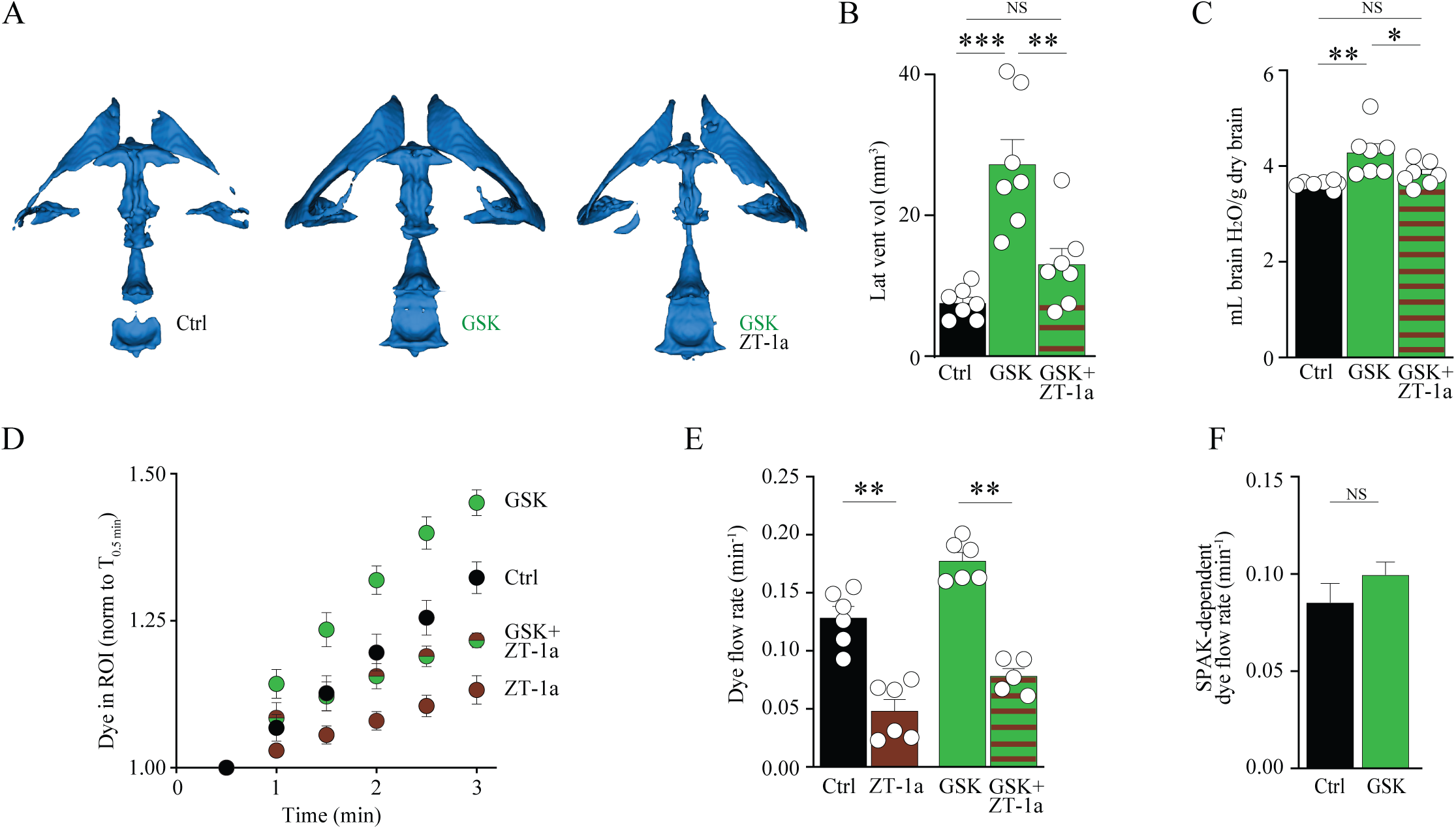
GSK101-induced CSF hyperdynamics depends on SPAK. **A** Representative 3D-volumetry of a control rat, a GSK101-treated rat, and a GSK101+ZT-1a-treated rat, demonstrating the ventricular volumes. **B** Lateral ventricle volumes of control rats (n = 7), GSK101-treated rats (n = 7), and GSK101+ZT-1a-treated rats (n = 7) quantified from MRI, see **Suppl. Fig. S5B** for representative MRI scans. **C** Brain water content quantified from control rats (n = 7), GSK101-treated rats (n = 7), and GSK101+ZT-1a-treated rats (n = 7). **E** Quantification of the dye intensity (as a proxy for CSF production) determined from linear regression in **D** over the 3-minute time window from control rats (n = 6), rats exposed to intraventricular GSK101 delivery (n = 6), intraperitoneal ZT-1a delivery (n = 6), or a combination of GSK101 and ZT-1a (n = 5). **F** SPAK-dependent dye flow rates upon GSK101-treatment. See Suppl. Fig. S1 for the collected data set. Statistical evaluation with *Student’s* t-test (panel F) or one-way ANOVA with Tukey’s post-hoc test (panel B, C, E). *P < 0.05, **P < 0.01, ***P < 0.001, NS: not significant.

To reveal if the TRPV4-mediated, SPAK-dependent modulation of CSF homeostasis is reflected in the CSF secretion rate, we determined the acute TRPV4-mediated increase in CSF secretion following systemic pre-treatment with the SPAK inhibitor ZT-1a. Inclusion of ZT-1a reduced the control CSF secretion rate from 0.13 ± 0.01 min^-1^, n = 6 to 0.05 ± 0.02 min^-1^, n = 6 upon ZT-1a treatment (P < 0.01, **Fig. 4D-E)**, and attenuated GSK-101-induced hypersecretion of CSF (from 0.18 ± 0.01 min^-1^ in GSK101-treated rats to 0.08 ± 0.01 min^-1^ in GSK101+ZT-1a-treated rats, n = 6 of each, P < 0.01, **Fig. 4D-E**). The SPAK-dependent fraction of the CSF secretion rate (0.08 ± 0.01 min^-1^, n = 6) was obtained as the difference between the CSF secretion rate in the control setting and that in the presence of ZT-1a and was similar to that obtained in the presence of GSK101 (0.10 ± 0.01 min^-1^, n = 6, P = 0.14, **Fig. 4F**). These data demonstrate that the TRPV4-mediated CSF hypersecretion requires SPAK activity.

## Discussion

The present study positions SPAK as a central regulator of CSF homeostasis and identifies the kinase as a mechanistic bridge between TRPV4 activation and the CSF-producing transport machinery of the choroid plexus. Our findings support a model in which NKCC1 and the Na⁺/K⁺-ATPase rely on constitutive SPAK activity to operate in concert at basal physiological conditions, but only NKCC1 activity being shaped by the upstream conductor TRPV4. This framework aligns with the emerging view that CSF production is dynamically tuned with a signalling transduction cascade spanning over multiple regulators, each of which could serve as future pharmacological targets to treat hydrocephalus and other pressure-related disorders [15, 22, 28–31].

A central observation demonstrates that systemic SPAK inhibition rapidly reduces CSF secretion in anesthetized rats via its suppression of Na⁺/K⁺-ATPase- and NKCC1-dependent activity, both of which contribute robustly to CSF secretion across the luminal membrane of the choroid plexus [24]. Notably, bumetanide-induced inhibition of NKCC1-mediated CSF secretion occurs solely with intracerebroventricular delivery, and not with i.p. administration [15, 21, 32]. This direct link between SPAK activity and these two key contributors to choroid plexus-mediated CSF secretion aligns with SPAK serving as a regulatory scaffold for these and other luminal ion transporters in rodent and porcine choroid plexus epithelial cells [14]. Our findings extend the established SPAK- NKCC1 axis in choroid plexus and other epithelia [14, 31, 33, 34] to include an additional layer of regulation involving SPAK-dependent activation of the Na⁺/K⁺-ATPase. With direct pharmacological targeting of NKCC1 or the Na⁺/K⁺-ATPase in hydrocephalus settings being constrained by systemic effects, neuronal modulation, and/or toxicity [35], an ability to modulate these transporters indirectly through an upstream kinase highlights SPAK as a physiologically relevant and more selective intervention point in conditions with disturbed CSF dynamics [14, 15, 31].

The clinical relevance of this pathway becomes evident in our rat model of brain hemorrhage. A mimicked hemorrhagic event induced the ventricular enlargement and brain water accumulation characteristic of posthemorrhagic hydrocephalus [15, 36–38]. Inhibition of SPAK with systemic delivery of the inhibitor two hours *after* the hemorrhagic insult mitigated the ventriculomegaly and reduced the total CSF space as well as the overall brain water content. These findings suggest that SPAK-dependent CSF hypersecretion is not merely a downstream correlate but a functional contributor to posthemorrhagic pathophysiology and positions SPAK as a promising future pharmaceutical target for hydrocephalus with potential to be targeted in a clinically relevant time window from the systemic face of the choroid plexus. Notably, SPAK-dependent regulation over time may preferentially be engaged under pathological conditions [15] characterized by heightened TRPV4 activity, inflammation, or compromised barriers [31]. Such patho-selectivity is advantageous for therapeutic development, as it implies that SPAK inhibition can attenuate pathological CSF hypersecretion without perturbing baseline CSF homeostasis.

Within the regulatory axis of SPAK-NKCC1-Na^+^/K^+^-ATPase, TRPV4 emerges as not only a central regulator of basal CSF secretion under physiological conditions [16], possibly via its mechanosensing modality [19] in analogy to that of Piezo 1 [39], but also as a potential context-dependent driver of CSF hypersecretion in pathological conditions [16, 22, 40]. The latter likely in a complementary fashion to that of the inflammation-induced TLR4-mediated SPAK activation [14, 15]. TRPV4 activation in healthy rats increased the CSF secretion rate and could, on its own, provoke a hydrocephalic condition and an overall increase in brain water content. This direct TRPV4-dependent modulation of the CSF dynamics is consistent with its involvement in hydrocephalus formation upon entry of the plasma lipid lysophosphatidic acid (LPA) in connection with brain hemorrhage [16] and with its inhibition attenuating ventriculomegaly in rats with either genetically- or hemorrhage-induced hydrocephalus [16, 22, 40]. Mechanistically, our data show that this TRPV4-mediated hypersecretory response is routed primarily through NKCC1: i) TRPV4 stimulation enhanced NKCC1 activity with no modulation of the Na⁺/K⁺-ATPase and ii) NKCC1, but not Na⁺/K⁺-ATPase, inhibition blunted the TRPV4-evoked CSF hypersecretion. Importantly, the TRPV4-mediated CSF hypersecretion and ventriculomegaly was abolished with SPAK inhibition, together indicating that TRPV4 relies on a SPAK-NKCC1 signalling axis to drive CSF hypersecretion. Constitutive activity of SPAK thus appears to suffice for its full activation of the Na^+^/K^+^-ATPase, whereas further SPAK activation may promote additional phospho-activation of NKCC1 in pathological settings, whether induced by TRPV4 activity (as here shown) and/or via complementary immune-related pathways [14, 15]. This finding demonstrates SPAK as the missing mechanistic link between TRPV4 activation by hemorrhage-derived mediators such as lysophosphatidic acid (LPA) [23] and the NKCC1 hyperactivation that contributes to ventriculomegaly associated with the acute phase of posthemorrhagic hydrocephalus in rats [16]. Future delineation of the time course of SPAK phosphorylation and activation in relation to the upstream activators and downstream NKCC1 activation would reveal the exact time course of these events leading up to the functional response.

Amongst the limitations of this study is the reliance on pharmacological inhibitors and activations. SPAK inhibition was obtained with ZT-1a, which is considered selective [31], although off-target effects cannot be fully excluded. Genetic or RNA-based approaches, such as conditional SPAK knockdown in the choroid plexus, or lipid nanoparticle-mediated co-delivery of SPAK siRNA [41] in our model, would strengthen causal inference. Moreover, the present work focuses on acute and subacute time windows; whether SPAK inhibition can modify chronic hydrocephalus or long-term neurological outcomes remains unknown, as does the potential translation to human patients.

In conclusion, we demonstrate that TRPV4 activation in healthy rats can promote hydrocephalus via its downstream activation of NKCC1, but not the Na^+^/K^+^-ATPase. SPAK appears not only to be a basal regulator of NKCC1 as well as the Na^+^/K^+^-ATPase, but also as a central integrator between TRPV4 activation and NKCC1 (hyper)activity. By demonstrating that SPAK inhibition, accordingly, dampens CSF secretion, attenuates TRPV4-induced hypersecretion, and ameliorates the acute phase of posthemorrhagic hydrocephalus development, this study highlights the TRPV4-SPAK axis as a promising therapeutic target for acute hydrocephalus and motivates further exploration of therapeutic windows, safety, long-term efficacy, complementarity to the TLR4 pathways, potential dependence of inflammatory agents, and future translation to the clinic.

## Materials and methods

### Animals

Experiments were conducted in 9–10 week-old male Sprague Dawley rats (Janvier Labs) that were housed in a temperature-controlled room with a 12 h:12 h light-dark cycle (6 am to 6 pm) with free access to a standard rodent pellet diet and tap water. All animal experimental work conformed to the legislation for animal protection and care in the European Community Council Directive (EU Directive 2010/63/EU) and was approved by the Danish Animal Experiments Inspectorate (license no. 2024-15-0201-01612 and 2021-15-0201-00867). Only male rats were used to eliminate the confounding effects of hormonal fluctuations associated with the estrous cycle on choroid plexus NKCC1 expression and thus potential CSF dynamics[42].

### Anaesthesia

The experimental animals were intraperitoneally injected with 6 mg ml^−1^ xylazine in sterile water as muscle relaxant (0.17 ml per 100 g body weight, pre-heated to 37 °C, ScanVet) followed by ketamine as anaesthetic (5 min post-xylazine administration; + 60 mg ml^−1^ ketamine in sterile water (0.17 ml per 100 g body weight, pre-heated to 37 °C, ScanVet). Isoflurane (Attane vet) was employed for the intraventricular hemorrhage procedure and magnetic resonance imaging (mixed with air and O2). 5% isoflurane was used to induce anaesthesia and 1.5–2.5% to sustain anaesthesia. The body temperature of the anesthetized rats was maintained at 37°C by a homeothermic monitoring system (Harvard Apparatus). For survival procedures, the rats were preoperatively given subcutaneous injections of the analgesic buprenorphine (0.4 mg kg^−1^, Sandoz, re-administered 12 h post-operatively) and carprofen (5 mg kg^−1^, Norbrook).

### Solutions

CSF dynamics experiments were conducted in HCO3^-^-containing artificial cerebrospinal fluid (aCSF; (in mM) 120 NaCl, 2.5 KCl, 2.5 CaCl2, 1.3 MgSO4, 1 NaH2PO4, 10 glucose, 25 NaHCO3, pH adjusted with 95% O2/5% CO2). In experiments where the CSF could not readily be equilibrated with 95% O2/5% CO2 (radioactive flux assays), we employed HEPES-buffered aCSF ((in mM) 120 NaCl, 2.5 KCl, 2.5 CaCl2, 1.3 MgSO4, 1 NaH2PO4, 10 glucose, 17 Na-HEPES, adjusted to pH 7.4 with NaOH). GSK10167A (TRPV4 activator, G0798, Sigma) was dissolved in DMSO (10 mM) and further diluted in HEPES-buffered aCSF for radioactive flux assays and western blotting (100 nM [16]), or HCO_3_ ^-^-containing aCSF for *in vivo* assays with intracerebroventricular delivery (5 µM [16]). Full solubility was verified with careful visual investigation. Of note, It has earlier been suggested that GSK10167A may be toxic to cultured choroid plexus epithelial cells [43]. However, *ex vivo* choroid plexus epithlieal integrity has been verified following 1h exposure to 500 nM GSK 10167A (Suppl. File in [16]) and the effects of GSK10167A are robustly attenuated in the present study upon inhibition of SPAK or NKCC1, suggesting the GSK101067A does not act via non-specific toxic side effects. ZT-1a (SPAK inhibitor, HY-136532, MedChemExpress) was dissolved in DMSO and further diluted in 0.9% NaCl for *in vivo* assays with intraperitoneal delivery (10 mg kg^-1^) [31] or in HEPES-buffered aCSF for radioactive flux assays (10 µM) [31]. For intraperitoneal administration, an equiosmolar NaCl solution with matched DMSO concentration served as vehicle. Ouabain (Na^+^/K^+^-ATPase inhibitor, Sigma, O3125) was dissolved directly into the HEPES-buffered aCSF for flux assays (2 mM [36]) or in HCO_3_ ^-^-containing aCSF for *in vivo* assays (2 mM intracerebroventricular) [24]. Bumetanide (NKCC1 inhibitor, Sigma, B302320) was dissolved in DMSO and further diluted in HEPES-buffered aCSF for flux assays (20 µM) or in HCO_3_ ^-^-containing aCSF for *in vivo* assays (200 µM intracerebroventricular) [24]. All solutions contained appropriate concentrations of vehicle (DMSO, D8418, Sigma), which amounted to 0.05–0.1% DMSO.

### Live CSF flow imaging

Anaesthetized rats were positioned in a stereotactic frame, with the cranium and upper neck muscles exposed. A burr hole was drilled 1.3 mm posterior and 1.8 mm lateral to the skull landmark bregma. A Hamilton syringe (RN 0.40, G27, a20, Agntho’s) was inserted 4 mm into the lateral ventricle. For drugs delivered intracerebroventricularly, the experiment was initiated with an injection of 15 μl of drug of interest (GSK101: 5 µM; ouabain: 2 mM; bumetanide: 200 µM) at a rate of 1.5 µl s^-1^. All pharmacological compounds are expected to be swiftly diluted (approximately 10-fold in the ∼150-200 μl total CSF pool) and thus reach both the ipsi- and contralateral ventricles upon ventricular entrance, which leads to estimated functional ventricular concentrations of 20 µM bumetanide, 200 µM ouabain [24], and 500 nM GSK101 [16]. The procedure was repeated after 5 min, but with carboxylate dye included (10 µM, IRDye 800CW, P/N 929-08972, LI-COR Biosciences), totalling a maximum of 10 min tissue exposure to the compounds. For intraperitoneal delivery, ZT-1a (10 mg kg^-1^) was administered 30 min pre-imaging. Image acquisition was initiated 1 min after carboxylate injection, continuing for 5 minutes with 30-second intervals using the Pearl Trilogy Small Animal Imaging System (LI-COR) (800 nm channel, 85 µm resolution). Rats were stabilized during imaging with a custom tooth holder. Fluorescence was measured over time at a region of interest (ROI) near the skull landmark lambda and quantified relative to the initial fluorescence intensity within the ROI at 30 seconds. After imaging, a white field image of the head was captured, followed by an examination of the isolated brain hemispheres to confirm bilateral carboxylate staining in the lateral ventricles. Data were analyzed using Image Studio 5.2 (LI-COR Biosciences, Nebraska, US). Animals were excluded (n = 3) if they did not respond to the initial anaesthesia regimen (to avoid ketamine re-dosing, which lowers the rate of CSF secretion [27]).

### Radioactive flux assays

Rats were anesthetized and euthanized by decapitation. The brains were immediately isolated and immersed in ice-cold HEPES-aCSF for 10 min, followed by separation of the two hemispheres and isolation of the choroid plexus from the lateral ventricles. The isolated lateral choroid plexus was allowed to recover for 10 min in 37°C HEPES-aCSF followed by incubation in an isotope solution containing rubidium (^86^Rb^+^) (1 µCi ml^-1^, 022-105724-01624-0001, POLATOM) and ^3^H-mannitol (4 µCi ml^-1^, NET101, Perkin Elmer). ^3^H-mannitol does not enter the choroid plexus epithelial cells and serves as an extracellular marker and ^86^Rb^+^ acts as a congener for K^+^ and therefore readily enters the cells via the ion transport mechanisms in choroid plexus [35]. For influx, the choroid plexus was treated with inhibitors/activators as indicated in the results section for 10 min before placement in the isotope solution containing the same inhibitors (2 min) and subsequently rinsed in ice-cold isotope-free HEPES-aCSF containing 2 mM ouabain, 20 µM bumetanide, and 100 µM BaCl2 (to prevent efflux of intracellular ^86^Rb^+^ during the washing procedure), followed by transfer into scintillation vials containing 200 µl Solvable (6NE9100, Perkin Elmer). For efflux experiments, the choroidal tissue accumulated ^86^Rb^+^ and ^3^H-mannitol for 10 min with the inclusion of inhibitors/activators as indicated in the results section before the tissue was swiftly rinsed in 37°C isotope-free HEPES-aCSF. The tissue was then transferred into new wells containing isotope-free HEPES-aCSF with inhibitors/activators as indicated in the results section, at 10 s intervals. For every time point, 200 µl of surrounding HEPES-aCSF was subsequently collected and placed into scintillation vials. At the end of the experiment, the choroid plexus was put into a scintillation vial containing 200 µl Solvable. For both influx and efflux, the choroid plexus was dissolved completely before the isotope content was determined in 2 ml Ultima Gold™ XR scintillation liquid (6013119, Perkin Elmer) using the Tri-Carb 2900TR Liquid Scintillation Analyzer (Packard). Efflux data are shown as the natural logarithm of the ^86^Rb^+^ activity at each time point [16] normalized to the initial ^86^Rb^+^ activity (A0) as a function of time. The slope from linear regression analysis was used to determine the ^86^Rb^+^ efflux rate constant (min^−1^) [35].

### Brain water content

Healthy rats were anaesthetised as for the IVH model, and a burr hole was drilled for the injection of GSK101 intracerebroventricularly (5 µM) or GSK101 + ZT-1a intraperitoneally (10 mg kg^-1^) or vehicle 24 h before sacrifice. Rats with induced IVH were treated with ZT-1a intraperitoneally 2 h and 12 h post-insult. Following surgery, the animals were allowed to recover in their home cage. 24 h post-surgery, the rats were anesthetized with a mix of 6 mg ml^−1^ xylazine and 60 mg ml^−1^ ketamine in sterile water (intraperitoneally, 0.17 ml per 100 g body weight, pre-heated to 37 °C, ScanVet), sacrificed, and the brain was quickly removed, placed in a pre-weighed porcelain beaker (Witeg), and weighed within 1 min of isolation. The brain tissue was dried at 100°C for 72 h to a constant mass. The dry brain was then weighed, and the brain water content was calculated in ml gram^-1^ dry weight using the formula: (wet weight - dry weight) / dry weight.

### Rat model of IVH

The surgery was performed on isoflurane-anesthetized rats under aseptic conditions, with body temperature maintained at 37°C using a rectal probe and feedback-controlled heating pad (Harvard Apparatus). Rats were placed in a stereotaxic frame (Harvard Apparatus), and a midline incision was made to expose the skull. A burr hole was drilled above the right lateral ventricle (0.6 mm posterior, 1.6 mm lateral to bregma). Blood was drawn from the tail artery using a 25-gauge needle, and immediately thereafter, 200 µl of autologous blood (or saline for sham operations) was injected into the right lateral ventricle (4.5 mm ventral) over 15 min using a 27-gauge needle and an automated 11 Pico Plus Elite mini pump (Harvard Apparatus) [44]. This blood volume will disperse throughout the ∼150 µl total CSF volume and promotes reproducable posthemorrhagic hydrocephalus in rats with no apparent macroscopic brain damage in the 24h experimental window [36, 45]. The needle remained in place for 5 min. to prevent backflow. The burr hole was plugged with Spongostan (Ethican) and covered with surgical mesh, the skin was sutured, and the rats were allowed to recover before being returned to their housing. 2 h post-insult, the rats were treated intraperitoneally with ZT-1a (10 mg kg^-1^) or vehicle (re-dosed after 12 h).

### Magnetic resonance imaging (MRI)

Anesthetized rats underwent MRI in a 9.4 Tesla preclinical horizontal bore scanner (BioSpec 94/30 USR, Bruker BioSpin) equipped with a 240 mT/m gradient coil (BGA-12 S, Bruker) at the Preclinical MRI Core Facility, University of Copenhagen. The scanner was interfaced to a Bruker Avance III console and controlled by Paravision 6.1 software (Bruker). Imaging was performed with an 86 mm inner-diameter volume resonator and a 4-channel surface quadrature array receiver coil. The animal body temperature was maintained at 37 ± 0.5 °C with a thermostatically controlled waterbed, and its respiratory rate was monitored by an MR-compatible monitoring system (SA Instruments). The imaging protocol consisted of T2-weighted 2D rapid acquisition with relaxation enhancement (2D-RARE) for reference spatial planning with the following settings: repetition time (TR) = 4000 ms, effective echo time (TE) = 60 ms, number of averaging [23] = 4, RareFactor = 4, slice thickness = 500 μm, in-plane resolution = 137 × 273 μm, 25 coronal slices, total acquisition time [23] = 8.5 minutes. For obtaining high resolution CSF volumetry, a 3D constructive interference steady-state sequence (3D-CISS) image was calculated as a maximum intensity projection (MIP) from 4 realigned 3D-TrueFISP volumes with 4 orthogonal phase encoding directions (TR = 4.6 ms, TE = 2.3 ms, NA = 1, Repetitions = 2, Flip angle = 50°, 3D spatial resolution 100 × 100 × 100 μm, RF phase advance 0, 180, 90, 270°, TA = 28 minutes). To obtain optimal spatial uniformity, all acquired 3D-TrueFISP volumes were motion-corrected before calculation as MIP, and the image bias field was removed with Advanced Normalization Tools (ANTs). For each brain sample, the total brain volume was automatically segmented by using region growing with ITK-snap (version 3.8.0). In addition, the pixel intensity factorized semi-automatic thresholding was performed to segment the lateral ventricle in each hemisphere. The volume measurement of the whole brain and lateral ventricles was performed in ITK-snap. The analysis was carried out in a blinded fashion after 24 h treatment with vehicle, GSK101 (5 µM, intracerebroventricularly) or GSK101+Zt-1a (5 µM intracerebroventricularly + 10 mg kg^-1^, intraperitoneal) in control or IVH animals, respectively.

### Western blotting

Rats were anesthetized and euthanized by decapitation. The brains were immediately isolated and immersed in ice-cold HEPES-aCSF for 10 minutes, followed by separation of the two hemispheres and isolation of the choroid plexus from the lateral ventricles. The isolated lateral choroid plexus was allowed to recover for 10 minutes in 37°C HEPES-aCSF. The choroid plexus was treated with either GSK101 (100 nM) for 90 sec or vehicle. The choroid plexus samples were subsequently lysed in RIPA buffer (in mM: 150 NaCl, 50 Tris pH 8.0, 5 EDTA and 0.5% sodium deoxycholate, 0.1% SDS and 1% Triton X-100) with 0.4 mM pefabloc, 8 µM leupeptin, and PhosStop, sonicated (twice; 70% power for 30 s, Sonopuls, Bandelin), and centrifuged for 5 minutes at 10.000 *g*. The samples were loaded on precast SDS-PAGE gels (4–20% Criterion TGX, Bio-rad) and immobilon-FL membranes (Merck Millipore) employed for the transfer. Primary and secondary antibodies were diluted 1:1 in Odyssey blocking buffer (LI-COR): PBS-T. Primary antibodies: anti-GAPDH (AB2302, Millipore, 1:5000), anti-NKCC1 (hpa020130, Sigma), anti-pNKCC1 (S763B, Dundee University). Secondary antibodies: IRDye 680RD donkey anti-chicken (LI-COR, P/N 925-68075, 1:10,000; targeting GAPDH), rabbit anti-sheep (Thermofisher, SA5-10060, 1:1000, targeting NKCC1) and donkey anti-rabbit (Invitrogen, A10043, 1:1000, targeting pNKCC1). Images were obtained by the Odyssey CLx imaging system and analyzed by Image Studio 5.2.5 (LI-COR).

### Data presentation and statistics

All data are presented as mean ± SEM and illustrated as scatter plots overlying histograms to illustrate biological variance, unless the few instances where two averages were deducted from eachother, in which case the data are shown as a histogram only. Statistical significance was tested with Student’s *t*-test or one-way ANOVA with Tukey’s multiple *post-hoc* test as indicated in the figure legends. P values < 0.05 were considered statistically significant. The number of experiments (*n*) corresponds to the number of independent animals. For western blotting, the two lateral choroid plexuses from each animal were split for pairwise comparison. For other experimental series (flux assays) the animals and choroid plexuses were randomily assigned to different conditions asuring that the same plexuses from individual animals were not both included in the same group. MRI segmentation was done in a blinded fashion. Live-imaging and radioisotope flux assays were performed for all groups in parallel (see **Suppl. Fig. S1**, and mentioned in relevant figure legends) but analysed according to each experimental question, as described in the Results section.

## Data availability

Datasets are available upon reasonable request.

## Author contributions

Conception and design of research: T.L.T-B. and N.M.; Conduction of the experiments: T.L.T-B. Analysis of data: T.L.T-B. Interpretation of results: T.L.T-B. and N.M.; Preparation of figures: T.L.T-B.; Drafting of the manuscript: T.L.T-B. and N.M.; Revision and approval of manuscript: T.L.T-B. and N.M.

## Acknowledgements

We are grateful for assistance from technician Anette D. Kaas.

---

**Fig S1.**
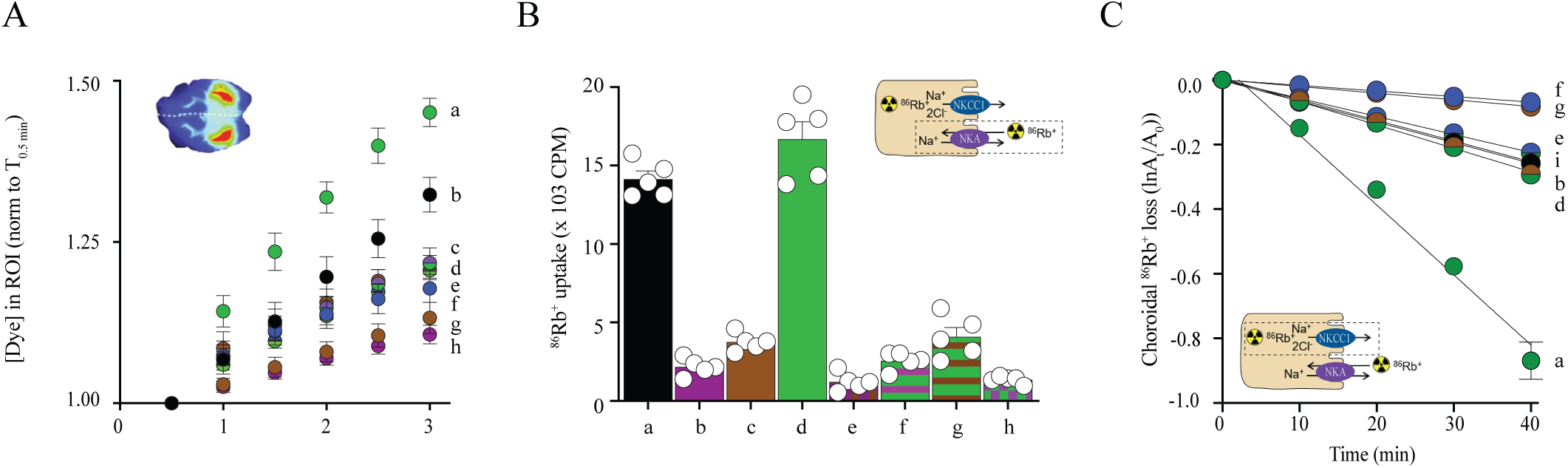
Live-imaging of dye intensity (as a proxy for CSF production) over a 3 min. time window (A) and radioisotope flux assays (B-C) with the inclusion of all employed drugs. ^86^Rb^+^ is a congener for K^+^ and was used to quantify the transport activity mediated by the Na^+^-K^+^-ATPase (B) and NKCC1 (C). **a**: GSK101; **b**: control; **c**: GSK101 + ouabain; **d**: GSK101 + ZT-1a; **e**: GSK101 + bumetanide; **f**: bumetanide; **g**: ZT-1a; **h**: ouabain; **i**: GSK101 + bumetanide + ZT-1a

**Fig S2.**
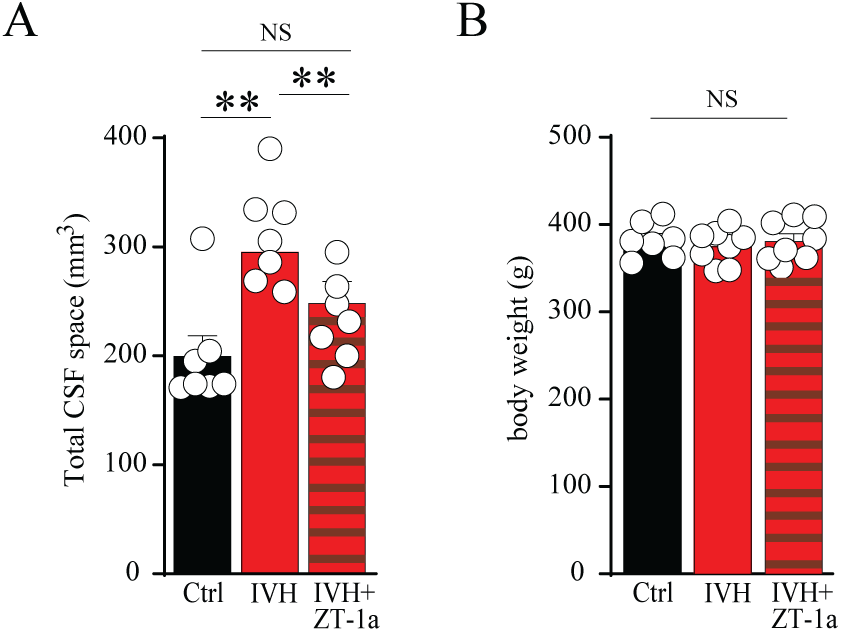
Quantification of total CSF spaces from MRI (A) and brain water (B) in control rats, rats with an intraventricular hemorrhage with or without ZT-1a treatment, n = 7 of each. Statistical evaluation with one-way ANOVA. **P < 0.01; NS: not significant.

**Fig S3.**
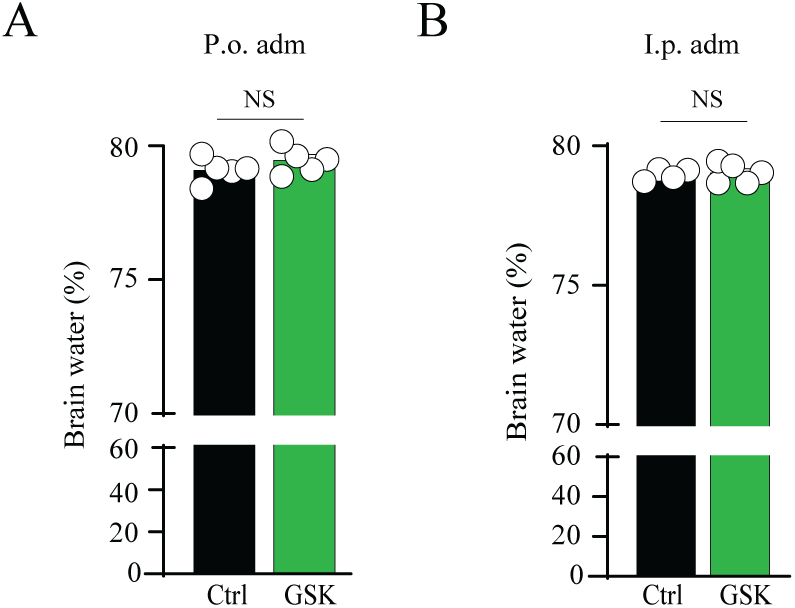
Non-systemically delivery of GSK101. Quantification of brain water in healthy control rats after GSK101 delivered per orally (A) or intraperitoneally (B), n = 4 - 5. Statistical evaluation with Student’s t-test. NS: not significant.

**Fig S4.**
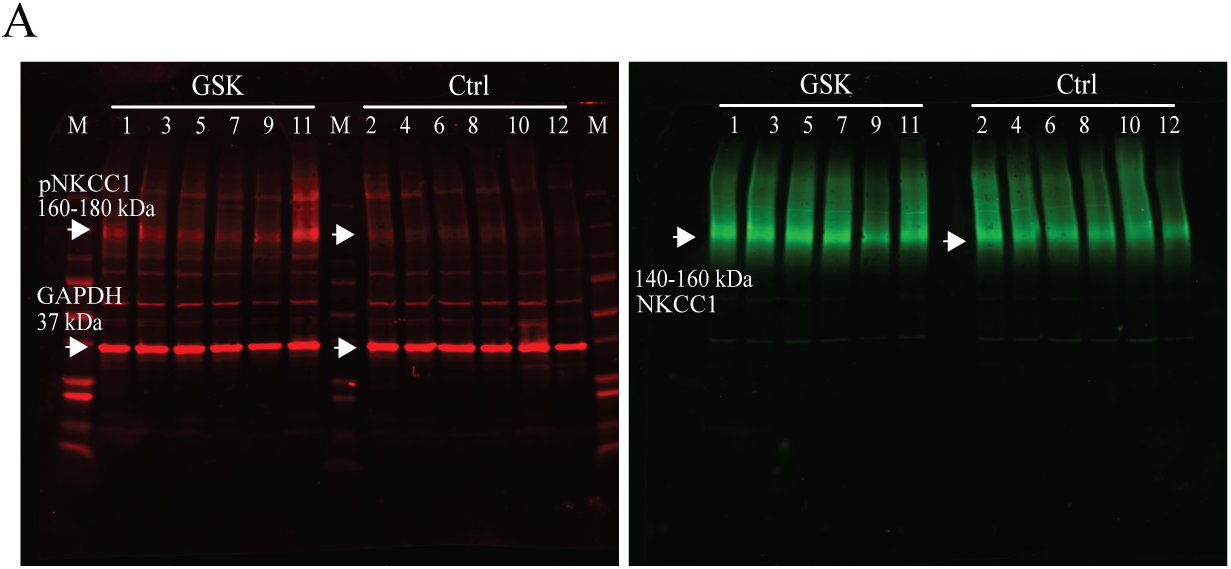
Western blotting of isolated lateral choroid plexuses from rats treated with vehicle (’Ctrl’) or GSK101 for 90 sec. The two plexuses from each rat were split into the non-treated (’Ctrl’) and GSK-group (1+2, 3+4 and so forth). Biological replicates: 6 rats. The membrane is shown as separated in 680nm and 800nm channel, respectively, and shows GAPDH as loading control, NKCC1 and phosphorylated NKCC1 (pNKCC1).

**Fig S5.**
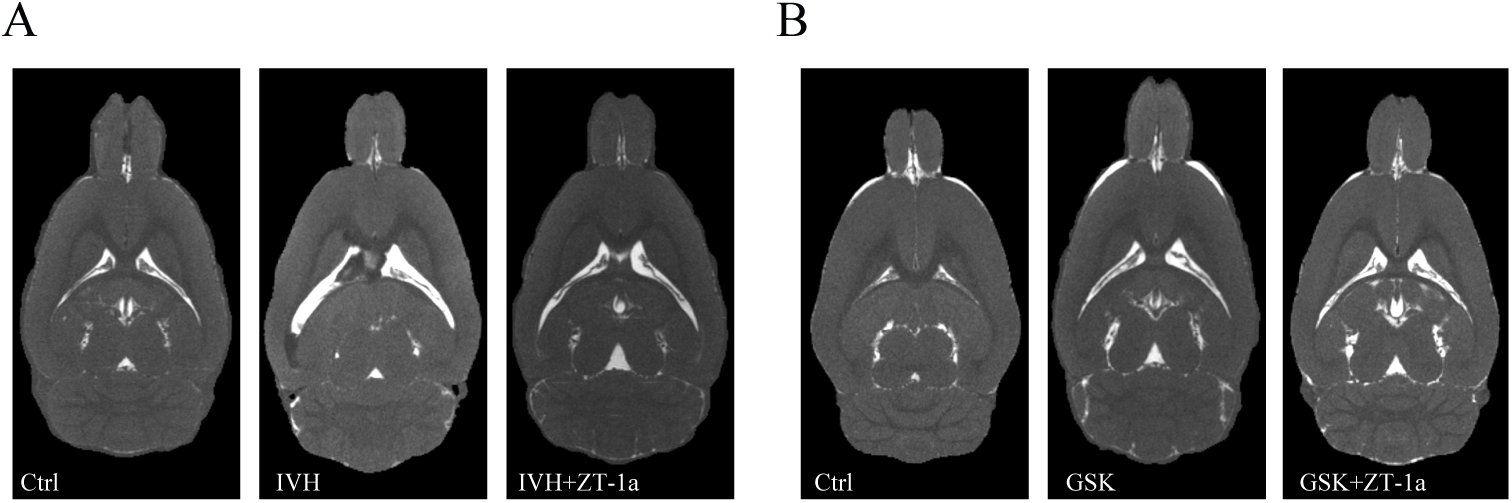
Representative T2 -weighted MRI rat brain sections 24 h after injection of saline (ctrl), autologous blood into the right lateral ventricle (IVH), and IVH with ZT-1a-treatment (i.p.) 2 h post-insult (IVH), n = 6-8 (A), and ctrl, GSK, or GSK with ZT-1a (i.p), n = 7 (B).

