## Supplementary material for "SPAK as a candidate mediator of TRPV4-induced NKCC1 activation and a possible pharmacological entry point in posthemorrhagic hydrocephalus"

### Supplementary file

## S1

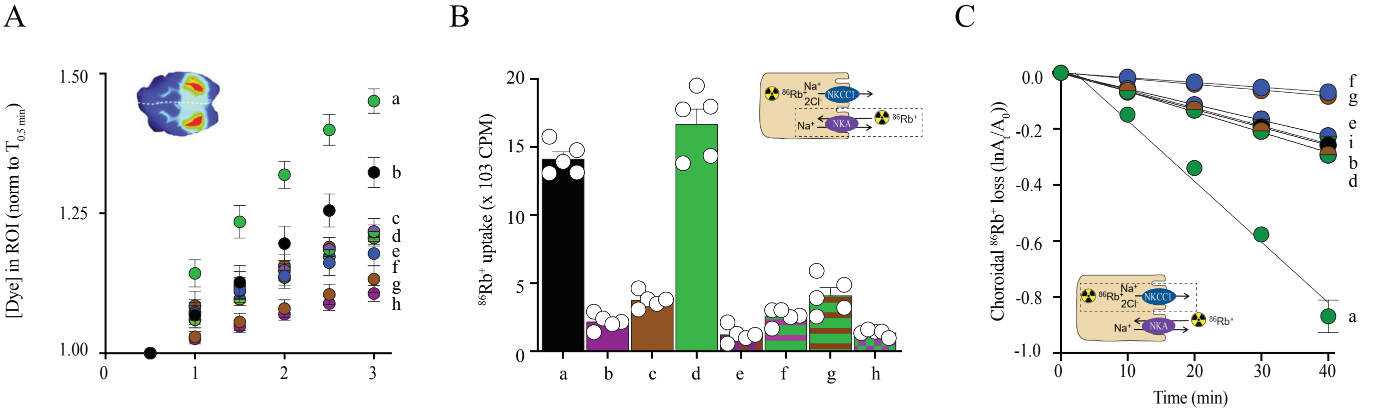


**S1** Live-imaging of dye intensity (as a proxy for CSF production) over a 3 min. time window (A) and radioisotope flux assays (B-C) with the inclusion of all employed drugs. $^{86}Rb^{+}$ is a congener for $K^{+}$ and was used to quantify the transport activity mediated by the $Na^{+}$-$K^{+}$-ATPase (B) and NKCC1 (C). **a**: GSK101; **b**: control; **c**: GSK101 + ouabain; **d**: GSK101 + ZT-1a; **e**: GSK101 + bumetanide; **f**: bumetanide; **g**: ZT-1a; **h**: ouabain; **i**: GSK101 + bumetanide + ZT-1a

## S2

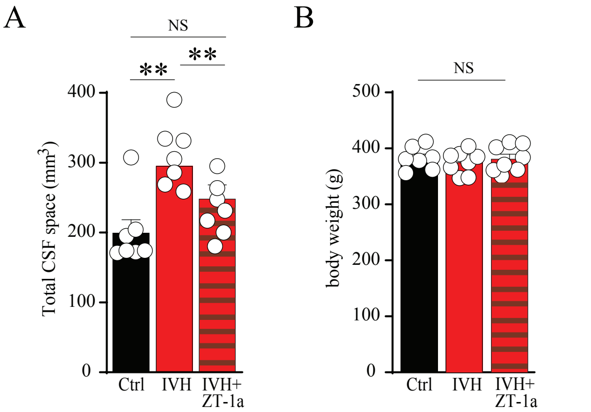


**S2** Quantification of total CSF spaces from MRI (A) and brain water (B) in control rats, rats with an intraventricular hemorrhage with or without ZT-1a treatment, n = 7 of each. Statistical evaluation with one-way ANOVA.
**$P < 0.01$; NS: not significant.

## S3

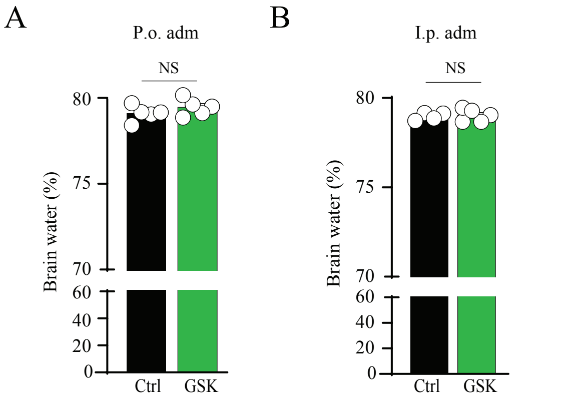


**S3** Non-systemically delivery of GSK101. Quantification of brain water in healthy control rats after GSK101 delivered per orally (A) or intraperitoneally (B), n = 4 - 5. Statistical evaluation with Student's t-test. NS: not significant.

## S4

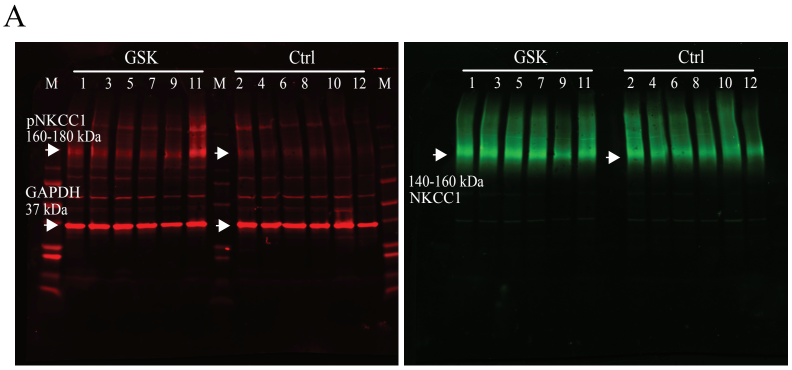


**S4** Western blotting of isolated lateral choroid plexuses from rats treated with vehicle ('Ctrl') or GSK101 for 90 sec. The two plexuses from each rat were split into the non-treated ('Ctrl') and GSK-group (1+2, 3+4 and so forth). Biological replicates: 6 rats. The membrane is shown as separated in 680nm and 800nm channel, respectively, and shows GAPDH as loading control, NKCC1 and phosphorylated NKCC1 (pNKCC1).

## S5

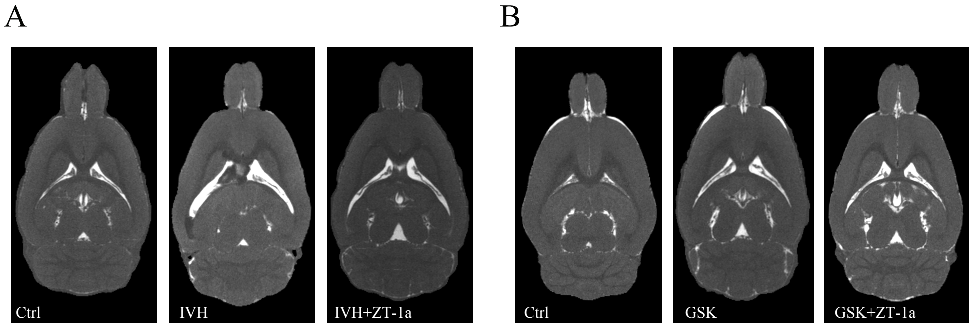


**S5** Representative T2 -weighted MRI rat brain sections 24 h after injection of saline (ctrl), autologous blood into the right lateral ventricle (IVH), and IVH with ZT-1a-treatment (i.p.) 2 h post-insult (IVH), n = 6-8 (A), and ctrl, GSK, or GSK with ZT-1a (i.p), n = 7 (B).
